# Identifiability and experimental design in perturbation studies

**DOI:** 10.1101/2020.02.03.931816

**Authors:** Torsten Gross, Nils Blüthgen

## Abstract

**Motivation:** A common strategy to infer and quantify interactions between components of a biological system is to deduce them from the network’s response to targeted perturbations. Such perturbation experiments are often challenging and costly. Therefore, optimising the experimental design is essential to achieve a meaningful characterisation of biological networks. However, it remains difficult to predict which combination of perturbations allows to infer specific interaction strengths in a given network topology. Yet, such a description of identifiability is necessary to select perturbations that maximize the number of inferable parameters.

**Results:** We show analytically that the identifiability of network parameters can be determined by an intuitive maximum flow problem. Furthermore, we used the theory of matroids to describe identifiability relationships between sets of parameters in order to build identifiable effective network models. Collectively, these results allowed to device strategies for an optimal design of the perturbation experiments. We benchmarked these strategies on a database of human pathways. Remarkably, full network identifiability was achieved with on average less than a third of the perturbations that are needed in a random experimental design. Moreover, we determined perturbation combinations that additionally decreased experimental effort compared to single-target perturbations. In summary, we provide a framework that allows to infer a maximal number of interaction strengths with a minimal number of perturbation experiments.

**Availability:** IdentiFlow is available at github.com/GrossTor/IdentiFlow.

## Introduction

Rapid technological progress in experimental techniques allows to quantify a multitude of cellular components in ever increasing level of detail. Yet, to gain a mechanistic understanding of the cell requires to map out causal relations between molecular entities. As causality cannot be inferred from observational data alone (Pearl, 2009), a common approach is to observe the system’s response to a set of localised perturbations (Sachs *et al.*, 2005) and reconstruct a directed interaction network from such data. A recurring idea within the large body of according network inference methods (Marbach *et al.*, 2010) is to conceive the system as ordinary differential equations and describe edges in the directed network by the entries of an inferred Jacobian matrix (Gardner *et al.*, 2003; Bonneau *et al.*, 2006; Tegner *et al.*, 2003; Kholodenko, 2007; Bruggeman *et al.*, 2002; Timme, 2007). Such methods have been successfully applied to describe various types of regulatory networks in different organisms (Ciofani *et al.*, 2012; Arrieta-Ortiz *et al.*, 2015; Lorenz *et al.*, 2009; Klinger *et al.*, 2013; Brandt *et al.*, 2019). They are continuously improved, e.g. to reduce the effect of noise, incorporate heterogeneous data sets, or allow for the analysis of single cell data (Greenfield *et al.*, 2013; Santra *et al.*, 2018; Klinger and Blüthgen, 2018; Santra *et al.*, 2013; Kang *et al.*, 2015; Dorel *et al.*, 2018) and have thus become a standard research tool. Nevertheless, identifiability (Hengl *et al.*, 2007; Godfrey and DiStefano, 1985) of the inferred network parameters within a specific perturbation setup has not yet been rigorously analysed, even though a limited number of practically feasible perturbations renders many systems underdetermined (De Smet and Marchal, 2010; Meinshausen *et al.*, 2016; Bonneau *et al.*, 2006). Some inference methods do apply different heuristics, such as network sparsity, to justify parameter regularisation (Gardner *et al.*, 2003; Bonneau *et al.*, 2006; Tegner *et al.*, 2003), or numerically analyse identifiability through an exploration of the parameter space using a profile likelihood approach (Raue *et al.*, 2009). Yet, neither approach provides a structural understanding on how parameter identifiability relates to network topology and the targets of the perturbations. However, such structural understanding is required to systematically define identifiable effective network models and to optimize the sequence of applied perturbations. The latter is of particular interest because perturbation experiments are often costly and laborious, which demands to determine the minimal set of perturbations that reveals a maximal number of network parameters. To address these challenges, this work derives analytical results that explain the identifiability of network parameters in terms of simple network properties which allow to optimize the experimental design.

## Methods

We consider a network of n interacting nodes whose abundances, *x*, evolve in time according to a set of (unknown) differential equations

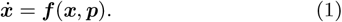

The network can be experimentally manipulated by p different types of perturbations, each represented by one of the p entries of parameter vector ***p***. We only consider binary perturbations that can either be switched on or off. Without loss of generality, we define ***f***(***x, p***) such that the *k*-th type of perturbation changes parameter *p*_*k*_ from its unperturbed state *p*_*k*_ = 0 to a perturbed state *p*_*k*_ = 1.

The main assumption is that after a perturbation the observed system relaxes into stable steady state, ***φ***(***p***), of Equation 1. Stability arises when the real parts of the eigenvalue of the n × n Jacobian matrix, ***J***_***ij***_ (***x, p***) = ***∂f***_***i***_(***x, p***)/***∂x***_***j***_, evaluated at these fixed points, ***x*** = ***φ***(***p***), are all negative within the experimentally accessible perturbation space (no bifurcation points). This implies that ***J*** (***φ***(***p***), ***p***) is invertible, for which case the implicit function theorem states that ***φ***(***p***) is unique and continuously dierentiable, and

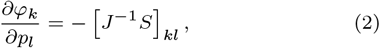

where n × p Sensitivity matrix entry, *S*_*ij*_ = *∂f*_*i*_(*x, p*)/*∂p*_*j*_, quantifies the effect of the *j*-th perturbation type on node *i*. Dropping functions’ arguments is shorthand for the evaluation at the unperturbed state, ***x*** = ***φ***(**0**) and ***p*** = **0**.

### A linear response approximation

A perturbation experiment consists of q perturbations, each of which involves a single or a combination of perturbation types, represented by binary vector ***p***, which forms the columns of the p × q design matrix *P*. The steady states after each perturbation, ***φ***(***p***), are measured and their differences to the unperturbed steady state form the columns of the n × q global response matrix *R*. Assuming that perturbations are sufficiently mild, the steady state function becomes nearly linear within the relevant parameter domain,

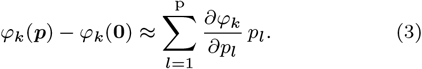

Replacing the partial derivative with the help of Equation 2 and writing the equation for all q perturbations yields

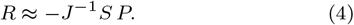

This equation relates the known experimental design matrix, *P*, and the measured global responses, *R*, to quantities that we wish to infer: the nodes’ interaction strengths, *J*, and their sensitivity to perturbations, *S*.

A dynamic system defined by rates ***f̃***(***x, p***) = ***W f***(***x, p***), with any full rank n × n matrix *W*, has the same steady states but different Jacobian and sensitivity matrices, namely *W J* and *W S*, as the original system, defined by Equation 1. It is thus impossible to uniquely infer *J* or *S* from observations of the global response alone, and prior knowledge in matrices *J* and *S* is required to further constrain the problem. In the following, we assume that prior knowledge exists about the network topology, i.e. about zero entries in *J*, as they correspond to non-existent edges. Likewise, we assume that the targets of the different types of perturbations are known, which implies known zero entries in *S* for non-targeted nodes. In line with prior studies (Kholodenko, 2007), we also fix the diagonal of the Jacobian matrix

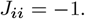

Thus, for the *i*-th row of *J* we can define index lists 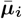 and 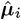 to identify its known and unknown entries. The first indicates missing edges or the self loop and the second edges going into node *i*. These lists have 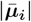 and 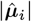 entries, respectively, with

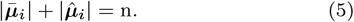

Analogously, for the *i*-th row of *S* we define index lists *v̅*_*i*_ and *v̂*_*i*_, with

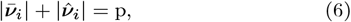

to report its unknown and known entries. These describe the perturbations that do not target or respectively target node *i*.

We show in Supplementary Material S1 that Equation 4 can be repartitioned to obtain a system of linear equations for each row in *J* and *S*, exclusively in the

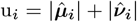

unknown parameters, which we collect in vector *x*_*i*_. Thus, there is a *u*_*i*_× *d*_*i*_ matrix *V*_*i*_, such that

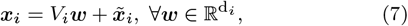

where *x̃*_*i*_ is some specific solution to the equation system. We further show in Supplementary Material S1 that *V*_*i*_ is a basis of the kernel of

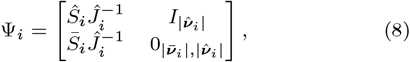

where 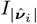and 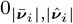 are the identity and zero matrix of annotated dimensionality. The 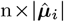 matrix 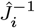 consists of the columns of (*J ^−^*^1^)^*T*^ that are selected by indices in *μ*_*i*_. Finally, |*v̅*_*i*_| × n matrix 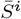 and *v̂*_*i*_ × n matrix 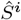 shall be formed by taking rows of *S*^*T*^ according to indices in *v̅*_*i*_ and *v̂*_*i*_. These matrix partitionings are demonstrated for a toy example in Supplementary Figure S1. Furthermore, in Supplementary Material S1 we derive the following expression for the solution space dimensionality

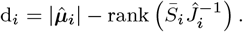

### Identifiability conditions

The system is underdetermined when *d*_*i*_ > 0. But independent of *d*_*i*_, a parameter is identifiable if the solution space is orthogonal to its according axis direction. This idea can be expressed as algebraic identifiability c onditions. A ccordingly, w e show in Supplementary Material S1 that the unknown interaction strength 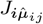 is identifiable if and only if

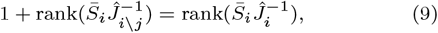

where 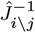 is matrix 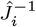 with the *j*-th column removed. Furthermore, the unknown sensitivity 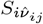 is identifiable if and only if

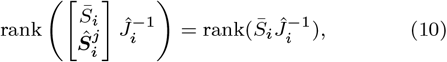

where 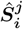 denotes the *j*-th row of matrix *Ŝ*_*i*_. However, the ranks depend on the unknown network parameters themselves and can thus not be directly computed. Yet, we can show how a reasonable assumption makes this possible and allows to express the identifiability conditions as an intuitive maximum flow problem.

First, we rewrite the identity *J*^−1^ *J* = *I*_*n*_ as

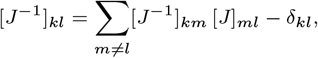

with *δ*_*kl*_ being the Kronecker delta (recall that *J*_*ll*_ = −1). We can view this equation as a recurrence relation and repeatedly replace the [*J*^−1^]_*km*_ terms in the sum. The sum contains non-vanishing terms for each edge that leaves node *l*. Therefore, each replacement leads to the next downstream node, so that eventually one arrives at

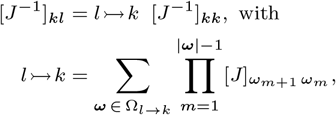

where the set *Ω*_*l→k*_ contains elements, *ω*, for every path from node *l* to node *k*, each of which lists the nodes along that path. Strictly speaking, these elements are walks rather than paths because some nodes will appear multiple times if loops exist between *l* and *k*. In fact, with loops, *Ω*_*l→k*_ contains an infinite number of walks of unbounded lengths. But as the real part of all eigenvalues of *J* are assumed negative, the associated products of interaction strengths converges to zero with increasing walk length.

To simplify our notation, we want to expand the network by considering perturbations ***v̅***_***i***_ as additional nodes, each with edges that are directed towards that perturbation’s targets. Furthermore, letting the interaction strength associated with these new edges be given by the appropriate entries in *S* we can rewrite the matrix product

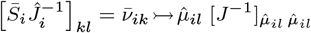

where 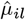 and *v̅*_*il*_ denote the *l*-th entry in 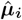 and ***v̅***_***i***_, respectively. As every finite-dimensional matrix has a rank decomposition, we can further write

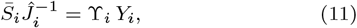

where 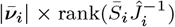 matrix *𝚼*_*i*_ and rank 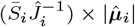 matrix *Y*_*i*_ have full rank. Finding such a decomposition therefore reveals the rank of 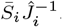. To this end, we propose

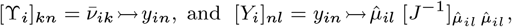

where *y*_*in*_ denotes the *n*-th component of a certain list of nodes where ***y***_***i***_. In order for Equation 11 to hold, it must be possible to split each path from any perturbation *v̅*_*il*_ to any node 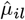 into a section that leads from the perturbation to a node in ***y***_***i***_ and a subsequent section that leads from this node to 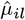. For an extended graph that includes an additional source node, with outgoing edges to each perturbation in ***v̅***_***i***_, and an additional sink node, with incoming edges from all nodes in 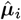 (see Figure 1B), ***y***_***i***_ thus constitutes a vertex cut whose removal disconnects the graph and separates the source and the sink node into distinct connected components. Next, we want to show that if ***y***_***i***_ is a minimum vertex cut, the rank of 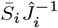 equals the size of ***y***_***i***_. Because Equation 11 is a rank decomposition this is equivalent to showing that the according matrices *𝚼*_*i*_ and ***y***_***i***_ have full rank. To do so, we apply Menger’s theorem (Menger, 1927), which states that the minimal size of ***y***_***i***_ equals the maximum number of vertex-disjoint paths from the source to the sink node. This also implies that each of these vertex-disjoint paths goes through a different node of the vertex cut ***y***_***i***_. Recall that entries in *𝚼*_*i*_ constitute sums over paths from perturbation to vertex cut nodes, so that we could write

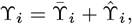

where 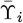 only contains the vertex-disjoint paths and 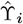 the sums over the remaining paths. As each of these vertex disjoint paths ends in a different vertex cut node, any column in 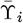 can contain no more than a single non-zero entry. Furthermore, as a consequence of Menger’s theorem there are exactly |***y***_***i***_| non-zero columns. Because these paths are indeed vertex disjoint also no row in 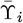 has more than a single non-zero entry. Thus, the non-zero columns are independent, showing that 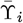 has full rank. The crucial assumption we want to make now is that the values of the interaction strengths lie outside a specific algebraic variety, which would render 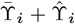 rank deficient. This would for example be the case if for a given vertex disjoint path there also is an alternative path whose associated product of interaction strengths has the same magnitude as that of the vertex disjoint path but opposite sign, making their sum vanish. This effect corresponds to a perfectly self-compensating perturbation. Most biological networks however cannot finetune their interactions to such a degree that they could achieve perfect self-compensation, which justifies this non-cancellation assumption.

**Figure 1:**
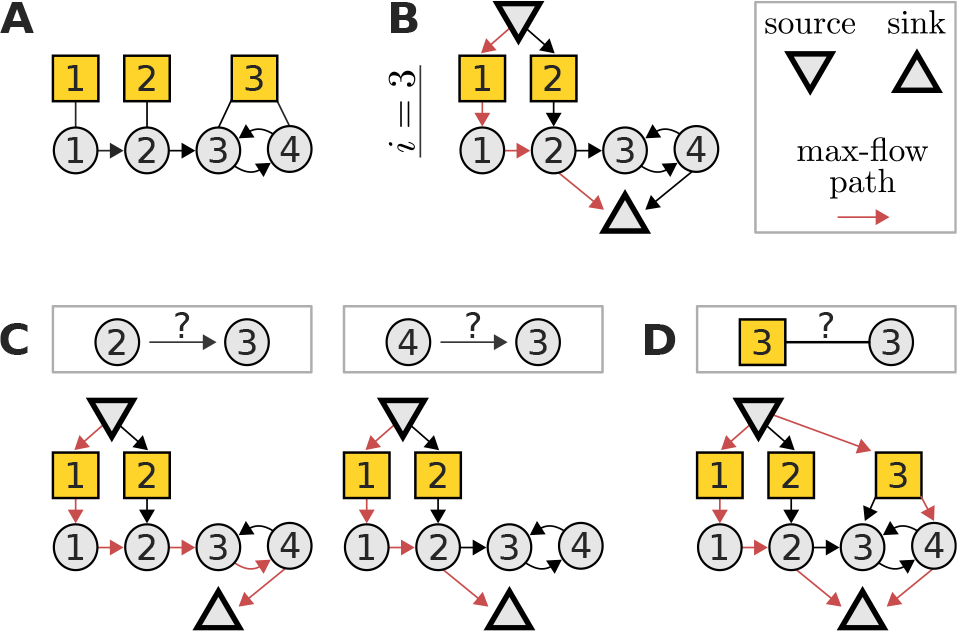
A maximum flow problem determines the identifiability of interaction strengths and perturbation sensitivities when reconstructing a network from perturbation data. (**A**) Example network with three perturbations (yellow squares) to illustrate the algorithm. (**B**) The corresponding flow network to determine the identifiability of the edges into node 3 and the sensitivity of node 3 to perturbations. A path carrying the maximal flow of one is denoted in red (note that it is not unique). (**C**) The interaction strength between a given node and node 3 is identifiable if and only if the maximum flow is reduced after removing that node’s edge to the sink node. In this example, there are alternative max-flow paths that re-establish a unit-flow after removal of the according edges. Thus, the respective interaction strengths are non-identifiable. (**D**) Similarly, the sensitivity of node 3 to perturbation 3 is identifiable, if and only if the depicted extension of the flow network does not increase the maximum flow. In this example, the maximum flow is increased by one, again revealing non-identifiability. Note that such flow representations provide an intuitive understanding on how alterations in the network or perturbation setting affect identifiability. For example, it is obvious that if the toy model would not contain an edge from node 3 to 4, the edge from 2 to 3 would become identifiable.

The same line of reasoning will demonstrate a full rank for matrix ***Y***_***i***_ as well, which implies that indeed

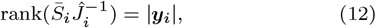

where ***y***_***i***_ is a minimum vertex cut between source and sink node. This equation has the crucial benefit that |***y***_***i***_| does not depend on any unknown parameters and can be computed as the maximum flow from source to sink node with all nodes having unit capacity (Ahuja *et al.*, 1993), as detailed in Figure 1B. This maximum flow problem can be solved in only 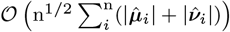, as shown in Theorem 6.3 in (Even and Even, 2012). More importantly though, it allows to express the algebraic identifiability conditions 9 and 10 in terms of network properties, providing an intuitive relationship between network topology, perturbation targets and identifiability. Specifically,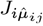 is identifiable if and only if the removal of the edge from node 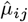 to the sink node reduces the maximum flow of the network, see Figure 1C, and 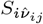 is identifiable if the maximum flow does not increase when an additional edges connects the source node with perturbation node *v̂*_*i,j*_ see Figure 1D.

### Identifiability relationships

Often, network inference is an underdetermined problem (De Smet and Marchal, 2010; Gross et al., 2019). Thus, to achieve identifiable effective network models, certain parameters have to be set to constant values, such that the remaining parameters become uniquely determinable. This requires an understanding of the identifiability relationships between parameters, i.e. we need to know which parameter becomes identifiable when other parameters are fixed. Supplementary Equation 16 formally relates these relationships to the ranks of certain linear subspaces of the range of 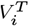 as defined in Equation 7. It shows that for each network node there is a set of parameters amongst which identifiability relationships can exist. Such a set contains those interaction strengths that quantify the edges, which target the associated node, and the associated node’s sensitivities to perturbations. Furthermore, we show in Supplementary Material S2 that the identifiability relationships of such parameter groups can be described as a matroid (Whitney, 1935). Matroids can be defined in terms of their circuits. Here, a circuit is a set of parameters with the property that any of its parameters becomes identifiable after fixing all of the others. Therefore, circuits describe all minimal parameter subsets that could be fixed to obtain an identifiable network.

We enumerated the set of circuits with an incremental polynomial-time algorithm (Boros et al., 2003). This algorithm requires an independence oracle that indicates linear dependence of subsets of columns of 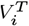. Supplementary Material S2 shows that we can construct such an oracle by considering linear dependence within the dual matroid, which amounts to determining

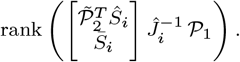

Matrices 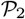 and 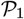 are truncated identity matrices defined in Supplementary Equations 16 and 17. Yet, the crucial point of this expression is that it has the same form as the left hand side of Equation 12. We can therefore conveniently determine it by solving a simple maximum-flow problem.

Supplementary Material S2 shows how to transform the circuits into cyclic flats. These provide a more convenient representation of the identifiability relationships, which we clarify at an example in the next section. Finally, certain scenarios constrain the choice of fixable parameters, for example when quantifying multiple isogenic cell lines (Bosdriesz *et al.*, 2018). Supplementary Material S2 describes a greedy algorithm that takes such preferences into consideration.

### Experimental design strategies

We assume that we are given a set of p perturbations, each of which targets a different subset of nodes. From these we want to select one or multiple perturbation sequences, according to a certain strategy, *s*. By means of our understanding of identifiability, we can determine *ξ*_*i*_, the number of identifiable edges after having performed the first *i* perturbations in such a sequence. Our goal is to find a strategy for which this number of identifiable edges increases fastest. Thus, as a measure of optimality of *s*, we can define an identifiability area under the curve

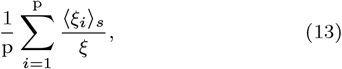

where 〈*ξ*_*i*_〉_*s*_ denotes the average of *ξ*_*i*_ over the ensemble of *s*-associated sequences, and *ξ* is the number of edges in the network. For any network and set of perturbations this score ranges between zero and one.

An obvious approach to design an optimal computational strategy is to simply consider the entire set of possible sequences and to select the ones that maximize the identifiability area under the curve. This is what we refer to as the *exhaustive* strategy. Clearly, it quickly becomes computationally intractable when the set of perturbations becomes large.

This is why we also propose strategies that build up the perturbation sequence in a stepwise manner and are therefore computationally efficient. At each step, the next perturbation is selected from a small set of perturbations that are deemed optimal according to a chosen strategy. Thus, identifying perturbation sequences in such a greedy optimisation approach amounts to performing a graph search over perturbation sets. We thus implemented a depth-first search to enumerate all strategy-conforming sequences. This sequence set might still be excessively large, when the number of possible perturbations is big. Therefore, we also implemented the option to randomly sample a fixed number of conforming sequences. A dynamic programming approach that avoids solving the same maximum-flow problem multiple times optimizes the search performance. Amongst the set of conforming strategies we eventually select the ones with the best performance according to Equation 13 (also see Figure 3A).

We suggest three different strategies to determine the set of the best next perturbations. The *single-target* approach selects those perturbations that minimize the overall dimensionality of the solution space, Σ_*i*_ *d*_*i*_, amongst the perturbations that maximize the number of identifiable edges. An alternative idea that does not make use of solution space considerations, is to assume that perturbations are the most informative when they cause a response at a maximal number of nodes. Accordingly, the *naive* strategy chooses perturbations that reach the largest number of nodes per perturbation target. Finally, we also want to allow for a combination of perturbations in a single experiment. Thus, we propose a *multi-target* strategy, which is similar to the single-target approach, except that it not only considers a single but any combination of perturbations. However, the entire power set of perturbation combinations might be too large to consider in practice. Therefore, we implemented an option for the multi-target strategy to build up combinations in a step-wise manner, where additional perturbations are only added to a combination if the enlarged combination increases performance. Additional details are provided in the documentation of the IdentiFlow package.

## Results

### Identifiability and identifiability relationships

Perturbation experiments are frequently used to infer and quantify interactions in biological networks. But whether a given network edge can indeed be uniquely quantified depends on the specific targets of the perturbations and the topology of the network. Yet in order to build interpretable network models and guide experimental design, we need to elucidate this identifiability status of the network parameters. Here, we view a biological system as a weighted directed network, and assume that perturbations are sufficiently mild to cause a linear steady state response. This allows to relate the interaction strengths between nodes (i.e. the entries in the Jacobian matrix *J*) and the sensitivity to perturbations (i.e. the entries in the Sensitivity matrix *S*) to the measured responses (Equation 4), an approach that is widely known as Modular Response Analysis (Kholodenko, 2007). We derived analytical identifiability conditions (Equations 9 and 10) that describe whether this relation allows to uniquely determine the network parameters for the given network topology and the experimental setting. However, these conditions can not be directly evaluated, as they depend on the (unknown) network parameters themselves. But instead, they can be reformulated as intuitive maximum flow problems, if one disregards singular conditions of self-cancelling perturbations.

The derivation and details are given in the Methods section but briefly, to determine the identifiability of either the interaction strength from node *j* to node *i*, or the sensitivity of node *i* to perturbation *p*, the following flow network is considered: The original network is extended by (i) adding a node for each perturbation that does not target node *i* and connecting it to the respective perturbation’s target(s), (ii) adding a “source” node that connects to all those perturbation nodes, and (iii) having all nodes that target node *i* connect to an additional “sink” node, see Figure 1B. Furthermore, all nodes (except source and sink) and all edges have a flow capacity of one. To reveal identifiability, we need to determine the network’s maximum flow from source to sink. This is a classic problem in computer science, which we solve using the Edmonds-Karp algorithm (Dinic, 1970; Edmonds and Karp, 1972) as implemented in the Networkx package (Hagberg *et al.*, 2008). Then, the interaction strength from node *j* to node *i* is identifiable if and only if the removal of the edge from node *j* to the sink node reduces the maximum flow, see Figure 1C. Similarly, node *i*’s sensitivity to perturbation *p* is identifiable if and only if the maximum flow does not increase after linking the source to an additional node that is in turn connected to all targets of perturbation *p*, see Figure 1D.

Often, experimental settings do not allow determining all un-known parameters (De Smet and Marchal, 2010; Gross *et al.*, 2019). Nevertheless, they constrain the solution space such that after fixing one or multiple parameters, others become identifiable. We found that such identifiability relationships can be described by matroids, which are combinatorial structures that generalize the notion of linear dependence (see Methods). This is demonstrated for an example perturbation experiment on the network displayed in Figure 2A.

**Figure 2:**
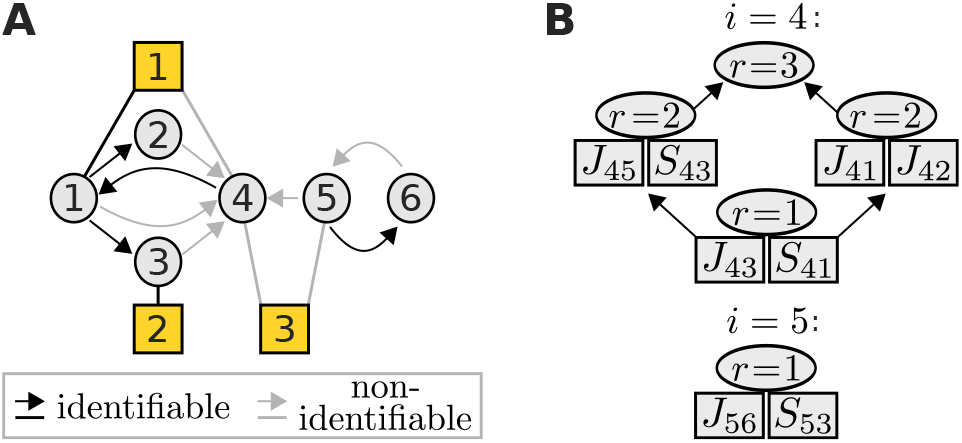
(**A**) An example network with three perturbations (yellow squares), where nodes 4 and 5 are associated with non-identifiable parameters (grey). (**B**) Their identifiability relationships are represented by the lattices of cyclic flats of rank *r*. Each cyclic flat consists of the annotated elements in addition to elements from its preceding cyclic flats. All parameters of a cyclic flat with rank *r* become identifiable if at least *r* independent flat parameters are fixed.

**Figure 3:**
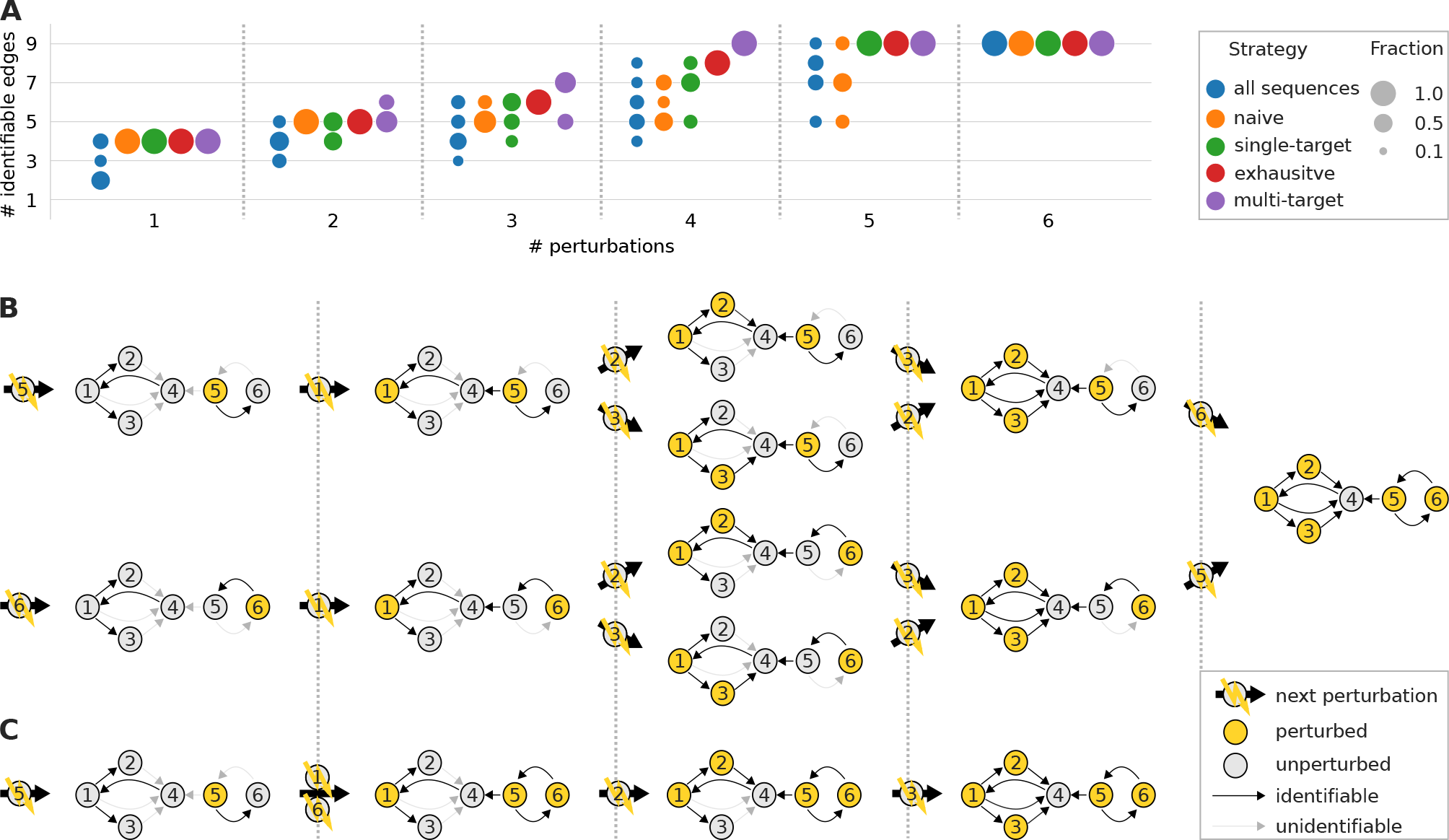
(**A**) The same network topology as in Figure 2 was subjected to a set of perturbations that target each node individually. Shown are distributions of numbers of identifiable edges for different experimental design strategies and an increasing number of perturbations. (**B**) All optimal (exhaustive) single-target perturbation sequences and (**C**) one multi-target sequence.

Each node is associated with a set of parameters amongst which identifiability relationships can exist. Such a set contains those interaction strengths, which quantify the edges that target the associated node, and that node’s sensitivities to perturbations. Here, nodes 4 and 5 are associated with sets of non-identifiable parameters. For example for node 5, these are *J*_56_ and *S*_53_. We represent the matroid for such a parameter set as a hierarchy (lattice) of cyclic flats, as show in Figure 2B. A cyclic flat is a set of parameters with an associated rank *r*. It has the property that all of its parameters become identifiable, if amongst them at least *r* independent parameters are fixed. Parameters are independent if none of them becomes identifiable after fixing the others. For node 5, parameters *J*_56_ and *S*_53_ only form a single cyclic flat with *r* = 1, and thus fixing either one parameter makes the other identifiable. The identifiability relationships among the six parameters associated with node 4 are more complex. For example, *J*_43_ and *S*_41_ form a cyclic flat with *r* = 1 and thus fixing one, fixes the other. Yet together with *J*_45_ and *S*_43_, they form a cyclic flat with *r* = 2, thus fixing e.g. *S*_41_ and *S*_43_ will allow unique determination of *J*_43_ and *J*_45_. In contrast, fixing *J*_43_ and *S*_41_ does not render any other parameter identifiable because they are not independent. This illustrates how the matroid description allows to generate effective models, i.e. models where a minimum number of parameters has to be set to fixed values to allow for a unique estimation of all other parameters. Importantly, the lattice of cyclic flats can be derived without specifying unknown parameters by solving a sequence of maximum flow problems (see Methods).

Collectively, our results provide a concise framework to algorithmically determine identifiability of network parameters and to construct identifiable effective networks when the experimental setting does not suffice to uniquely determine the original network structure.

### Experimental design

Next, we applied our identifiability analysis to optimize experimental design, i.e. to minimize the number of perturbation experiments that is required to uniquely determine a network’s interaction strengths. For this, we designed the following strategies to determine an optimal sequence from a set of available perturbations: The *exhaustive* strategy considers all possible sequences and selects the best performing amongst them. As this approach entails a prohibitive computational effort for larger networks, we also designed approaches that select perturbation sequences in a step-wise manner: First, the *single-target* strategy chooses the next perturbation such that it increases the number of identifiable edges most. Second, the *multi-target* strategy is similar to the single-target strategy except that it not only considers a single but any combination of perturbations. We then compared these strategies to a *naive* strategy that does not use our identifiability analysis. Rather, it chooses perturbations first that cause a response at a larger number of nodes (see Methods for details).

We first scrutinised the proposed experimental design strategies on the example network shown in Figure 2. We defined six different types of perturbations, each of which targets a (different) single node, or any combination of such for the multi-target strategy. Figure 3A shows how the number of identifiable edges increases with the number of performed perturbations for each strategy. A single strategy might propose multiple sequences, as described in Methods. Accordingly, Figure 3A shows the performance distribution over all such conforming sequences. In practice, we would only select the best performing sequence amongst them. Nevertheless, the depicted distributions are informative because for larger networks we can no longer enumerate all but only a (random) subset of conforming sequences.

When comparing the methods, we found that on average all strategies outperform randomly chosen sequences (“all sequences” distribution). Moreover, the “naive” strategy that did not use our framework mostly required all six perturbations to fully identify all parameters, whereas the single-target and exhaustive strategies only needed five, and the multi-target strategy only four perturbations. Figure 3B and Figure 3C display all perturbation sequences conforming to the single-target strategy, and one sequence conforming to the multi-target strategy respectively, and illustrate which network edge becomes identifiable at which step in the sequence.

To systematically analyse if and how our approach improves experimental design, we benchmarked the different strategies on all 267 nontrivial human KEGG (Kanehisa *et al.*, 2019) pathways, ranging from 5 to 120 nodes (see Supplementary Material S3 for details). Again, we assumed that perturbations can target (all) single nodes. For each network, we sampled 10 conforming sequences per strategy and compared against the performance of 10 randomly chosen sequences. As a performance measure, we considered the number of identified parameters as a function of the number of perturbations and computed a normalised area under the curve, as defined in Equation 13. Figure 4A shows the result of this benchmark, and confirms the trend already observed for the example in Figure 3A: Compared to choosing perturbations randomly, the naive strategy improved identifiability. Performance was further increased when we applied our single-target strategy, yet the multi-target strategy clearly performed best. An exhaustive enumeration of all sequences is not feasible for all KEGG networks. However, we found for a subset of small networks that there is no performance difference between the exhaustive and the single-target strategy, as shown in Supplementary Figure S4A.

**Figure 4:**
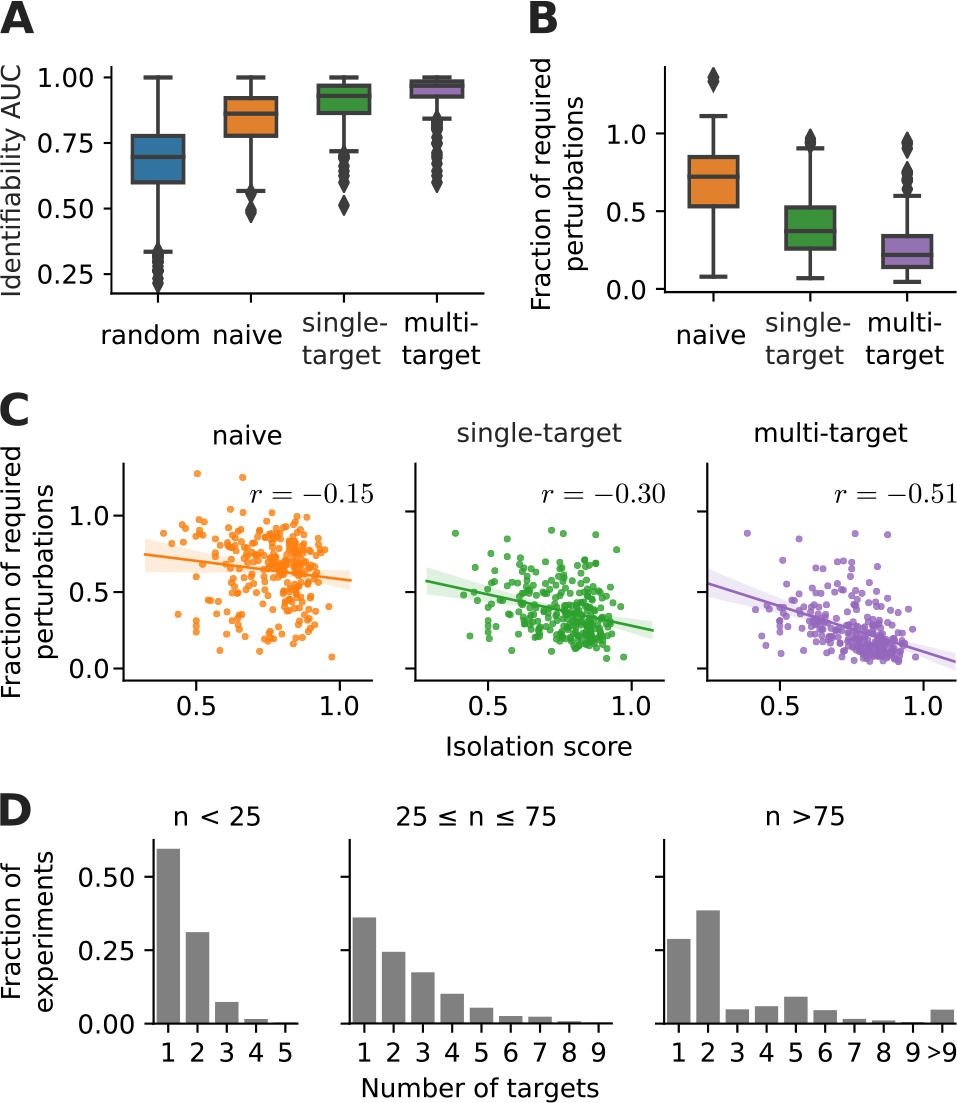
Performance of different experimental design strategies on 267 human KEGG pathways. (**A**) Identifiability AUC, defined as area under the number of identified nodes vs. number of perturbation curve, see Eqn. 13, (**B**) For each network and strategy, the average number of perturbations required for full identifiability is shown relative to the average number required for a random strategy. (**C**) The fraction of required perturbations correlated against the isolation score of a network (Eqn. 14), *r* : Spearman’s rank correlation. (**D**) The fraction of multi-target perturbations with a specific number of targets to all multi-target perturbations (experiments) in KEGG networks of the annotated size range.

Furthermore, we determined the average number of perturbations that is required for full network identifiability and computed the fraction between a given strategy and the random sequences, see Figure 4B. We found that the average number of required perturbations can be reduced to less than one third or even less than a quarter, when using a single-target or multi-target strategy, respectively.

We next investigated which network properties led to a performance increase using our strategies. Intuitively, perturbations might be more informative if their response propagates to large parts of the network. We therefore hypothesised that a careful experimental design is particularly beneficial when networks contain many isolated nodes with little connection to the rest of the network because, in contrast to a random choice, a good strategy could then avoid perturbing such non-informative targets. On the contrary, the sequence of perturbations is irrelevant in the extreme case of a fully connected network. To investigate this hypothesis we defined a network’s isolation score as

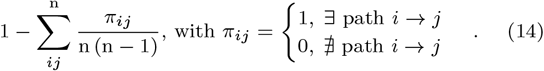

Figure 4C shows that indeed the isolation score negatively correlates with the previously defined fraction of perturbations required for full network identifiability. Furthermore, we also observed a positive correlation between isolation score and the difference in the identifiability AUC between non-random and random strategies, as shown in Supplementary Figure S4B. This suggests that indeed our experimental design strategies increase their performance with increasing network isolation.

When response signals converge at a node, the individual contribution from each incoming edge can not be distinguished. Thus, the advantage of a multi-target perturbation to potentially track signal propagation through larger parts of the network is counter-balanced if it leads to more convergent signal propagation. This is prevented when the (combined) perturbations target isolated parts of the network. Therefore the strongest correlation in Figure 4C is found for the multi-target strategy because with higher isolation score we can expect to find more such isolated subnetworks. And indeed, Figure 4D shows that the multi-target strategy typically suggest combinations of multiple single target perturbations, especially in larger networks.

In summary, we have developed an algorithmic approach to determine structural identifiability for a given network. This approach allows to derive experimental design strategies that drastically reduce experimental effort in perturbation studies. In particular, the multi-target strategy proved most efficient. Potentially, this finding has practical relevance because in many experimental contexts it easy to combine perturbations, e.g. by pooling ligands or inhibitors.

## Discussion

We have shown analytically that parameter identifiability in linear perturbation networks can be described as a simple maximum flow problem (summarised in Figure 1). This intuitive result not only explains how to achieve fully identifiable effective network models (Figure 2), but also enables us to optimize the design of perturbation experiments (Figure 3). As a test case, we examined all human KEGG pathways and found that our method typically allows to cut down the number of perturbations required for full identifiability to one fourth compared to choosing perturbation targets randomly (Figure 4). We provide a python implementation of our results github.com/GrossTor/IdentiFlow, which allows to determine identifiability, perform matroid computations that display identifiability relationships between parameters, and optimize experimental design. The package relies on standard maximum flow algorithms from the `Networkx` package (Hagberg *et al.*, 2008).

Technically, it would be possible to cope with non-identifiabilities numerically, as was done previously (Gardner *et al.*, 2003; Bonneau *et al.*, 2006; Tegner *et al.*, 2003; Dorel *et al.*, 2018) or even through the analysis of example networks. For the latter, we could set unknown parameters to random values and numerically compute the according ranks in the identifiability conditions Equation 9 and Equation 10. The idea is that a random example system is representative of all systems with the same topology and perturbation set-up. This approach would require to define certain thresholds to detect rank deficiency and the validity of the non-cancellation assumption. Even though it is therefore not guaranteed to work in general, we would still expect it to correctly determine a parameter’s identifiability in most cases. However, the crucial benefit of the maximum flow perspective is that identifiability can be intuitively understood in relation to the network topology and the targets of the perturbations. This means that instead of requiring numerical procedures on a case by case basis, we can directly see how the maximum flow depends on the network topology and perturbation set-up. This provides a comprehensive overview on which edges become identifiable under which perturbations. For one, this permits a straightforward optimisation of the experimental design, as shown before. But even in a situation where the set of perturbations is *a priori* fixed because of experimental constraints, our approach concisely reveals which network topologies are in principle amenable to a meaningful analysis. Thereby, it maps out the range of biological questions that are actually answerable with a given experimental set-up.

It is central to our analysis to assume mild perturbations that induce a linear steady state response. This also implies that the rates in Equation 1 depend linearly on the magnitude of the other nodes and the perturbations, as shown in Supplementary Material S1. But clearly, biological systems generally break linearity assumptions in varying degrees, which bears asking how useful our description is. In principle, we could expand the steady state function Equation 3 to higher orders and attempt to also infer nonlinear rate terms, which are products of different node and perturbation magnitudes. However such products no longer have any meaningful network interpretation, as they cannot be reasonably assigned to any edge. Therefore we argue that the linearity assumption is essential to derive a useful effective network description, if we choose to interpret the biological systems in terms of ordinary differential equations. Thus the identifiability analysis presented here stays valid even when the underlying system is highly nonlinear. On the downside, the biological meaning of interaction strengths becomes increasingly obscure the more the system violates the linearity assumption (Prabakaran *et al.*, 2014). Also, our assertions no longer hold if nonlinearities violate the non-cancellation assumption. Especially combinations of perturbations, as suggested by the multi-target experimental design strategy, might push the system into saturation and thus break our analysis. We therefore need to carefully consider such biological constraints as well.

Finally, we want to stress that our analysis solely describes structural identifiability. In contrast, practical identifiability concerns situations where parameters cannot be adequately determined because of a limited amount or a poor quality of experimental data (Raue *et al.*, 2011). This means that even when the identifiability condition for a specific parameter holds, it does not necessarily mean that its value can be reliably estimated. The maximum flow approach is currently agnostic to information about noise that could potentially render a structurally identifiable parameter practically non-identifiable. Similarly, it cannot handle situations where a measurement of a node’s steady state response is not only noisy but entirely missing. As this is a common challenge in novel single cell perturbation studies (Jaitin *et al.*, 2016; Datlinger *et al.*, 2017), we consider an according generalisation of our analysis as an important line of future research.

