## Supplementary Material for "Identifiability and experimental design in perturbation studies"

Sections S1 and S2 constitute an extended version of the sections on identifiability and identifiability relationships in the Methods part of the main text. They include additional information and examples, and explicit derivations that were abbreviated in the main text.

Section S3 provides additional details and data for the optimization of experimental design in KEGG pathways.

### S1 Identifiability

We want to consider a network of n interacting nodes whose abundances or magnitudes, $\boldsymbol{x}$, evolve in time according to a set of (unknown) differential equations

$$\dot{\boldsymbol{x}} = \boldsymbol{f}(\boldsymbol{x}, \boldsymbol{p}). \tag{1}$$

We assume that we can experimentally manipulate the system with p different types of perturbations, each of which is represented by one of the p entries of parameter vector $\boldsymbol{p}$. We shall only consider binary perturbations that can either be fully switched on or off. To keep notation simple and without loss of generality, we thus define $\boldsymbol{f}(\boldsymbol{x}, \boldsymbol{p})$, such that the $k$-th type of perturbation changes parameter $p_k$ from its unperturbed state $p_k = 0$ to a perturbed state $p_k = 1$.

One of the main assumptions about the observed system is that its temporal dynamics eventually relaxes into different constant states depending on the performed perturbation. These states are thought to represent stable fixed points, $\boldsymbol{\varphi}(\boldsymbol{p})$, of Equation 1, where stability arises because the real parts of the eigenvalue of the n × n Jacobian matrix, $J_{ij}(\boldsymbol{x}, \boldsymbol{p}) = \partial f_i(\boldsymbol{x}, \boldsymbol{p})/\partial x_j$, evaluated at these fixed points, $\boldsymbol{x} = \boldsymbol{\varphi}(\boldsymbol{p})$, are all negative within the experimentally accessible perturbation space (no bifurcation points). This implies that $J(\boldsymbol{\varphi}(\boldsymbol{p}), \boldsymbol{p})$ is invertible, for which case the implicit function theorem states that $\boldsymbol{\varphi}(\boldsymbol{p})$ is unique and continuously differentiable, and

$$\frac{\partial \varphi_k}{\partial p_l} = -\left[J^{-1} S\right]_{kl}, \tag{2}$$

where n×p sensitivity matrix entry, $S_{ij} = \partial f_i(\boldsymbol{x}, \boldsymbol{p})/\partial p_j$, quantifies the effect of the $j$-th perturbation type on node $i$. Dropping functions' arguments is shorthand for the evaluation at the unperturbed state, $\boldsymbol{x} = \boldsymbol{\varphi}(\boldsymbol{0})$ and $\boldsymbol{p} = \boldsymbol{0}$.

#### A linear response approximation

A perturbation experiment consists of q perturbations, each of which involves a single or a combination of perturbation types, represented by binary vector $\boldsymbol{p}$. These vectors shall form the p × q design matrix $P$. After each perturbation the system is allowed sufficient time until the newly established steady states, $\boldsymbol{\varphi}(\boldsymbol{p})$, can be measured. Let their differences to the unperturbed steady state form the columns of the n × q global response matrix $R$. The central approximation is to assume that perturbations are sufficiently mild, such that the steady state function becomes nearly linear within the relevant parameter domain,

$$\varphi_k(\boldsymbol{p}) - \varphi_k(\boldsymbol{0}) \approx \sum_{l=1}^{\mathrm{p}} \frac{\partial \varphi_k}{\partial p_l} p_l. \tag{3}$$

Replacing the partial derivative with the help of Equation 2 and writing the equation for all q perturbations yields

$$R \approx -J^{-1} S\, P. \tag{4}$$

Note that this equation holds exactly and independent of perturbation strength for a linear system

$$\dot{\boldsymbol{x}} = J\boldsymbol{x} + S\boldsymbol{p},$$

which can be seen by considering its steady state

$$\boldsymbol{x}^{\boldsymbol{0}} = J^{-1} S\, \boldsymbol{p}.$$

The crux of Equation 4 is that it relates the known experimental design matrix, $P$, and the measured global responses, $R$, to quantities that we wish to infer, namely the nodes' interaction strengths, $J$, and their sensitivity to perturbations, $S$. Thus, as a next step we shall rewrite the equation to disentangle the known and the unknown entries.

A dynamic system defined by rates $\tilde{\boldsymbol{f}}(\boldsymbol{x}, \boldsymbol{p}) = W\, \boldsymbol{f}(\boldsymbol{x}, \boldsymbol{p})$, with any full rank n × n matrix $W$, has the same steady states but different Jacobian and sensitivity matrices, namely $W\, J$ and $W\, S$, as the original system, defined by Equation 1. It is thus impossible to uniquely infer $J$ or $S$ from observations of the global response alone. However, some entries in matrices $J$ and $S$ might be known a priori and thus further constrain the problem. This is the case, when e.g. certain reactions rates are known. Typically however, such values are hard to come by. Rather, we assume prior knowledge about the network topology. That is, we know the zero entries in $J$ as they correspond to non-existent edges. Likewise, we assume to know the targets of the different types of perturbations which imply zero entries in $S$-rows corresponding to perturbations that that are known to not directly affect the network node associated with that row. In line with prior studies, we fix the diagonal of the Jacobian matrix

$$J_{ii} = -1.$$

Thus, for the $i$-th row of $J$ we can define index lists $\bar{\boldsymbol{\mu}}_i$ and $\hat{\boldsymbol{\mu}}_i$, with

$$|\bar{\boldsymbol{\mu}}_i| + |\hat{\boldsymbol{\mu}}_i| = \mathrm{n}, \tag{5}$$

identifying its known and unknown entries. The first correspond to missing edges or the self loop and the second to edges going into node $i$. Analogously, for the $i$-th row of $S$ we define index lists $\bar{\boldsymbol{\nu}}_i$ and $\hat{\boldsymbol{\nu}}_i$, with

$$|\bar{\boldsymbol{\nu}}_i| + |\hat{\boldsymbol{\nu}}_i| = \mathrm{p}, \tag{6}$$

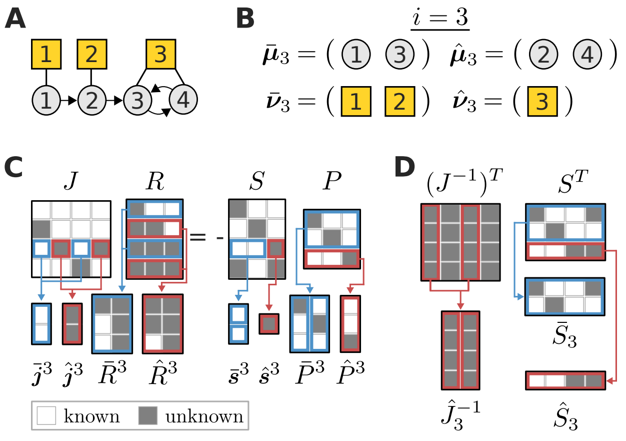


**Figure S1:** Three perturbations (yellow squares) are performed on a toy network (**A**). Network topology and perturbation targets determine the index lists from Equation 5 and Equation 6. Here they are depicted for $i = 3$ (**B**). A graphical representation of Equation 4 demonstrates the definition of various matrix partitions (**C** and **D**).

to report its unknown and known entries. These describe the perturbations that do not target or respectively target node $i$, see Figure S1B.

To see whether prior knowledge about $J$ and $S$ entries could render other entries determinable, we first rewrite Equation 4 as n linear equation systems

$$R^T \boldsymbol{j}_i = -P^T \boldsymbol{s}_i, \quad i = 1, 2, \ldots \mathrm{n}, \tag{7}$$

one for each column in $J^T$ and $S^T$, denoted as $\boldsymbol{j}_i$ and $\boldsymbol{s}_i$. Then, we collect the known and unknown $\boldsymbol{j}_i$-entries into vectors $\bar{\boldsymbol{j}}_i$ and $\hat{\boldsymbol{j}}_i$ following the indexing by $\bar{\boldsymbol{\mu}}_i$ and $\hat{\boldsymbol{\mu}}_i$. In the same manner, $\boldsymbol{s}^i$ is split into the known vector $\bar{\boldsymbol{s}}^i$ and unknown vector $\hat{\boldsymbol{s}}^i$ according to $\bar{\boldsymbol{\nu}}_i$ and $\hat{\boldsymbol{\nu}}_i$. To rewrite Equation 7 as a linear system of the unknown variables, we first partition its terms into known and unknown parts

$$R^T \boldsymbol{j}_i = \bar{R}_i\, \bar{\boldsymbol{j}}_i + \hat{R}_i\, \hat{\boldsymbol{j}}_i \quad \text{and} \quad P^T \boldsymbol{s}_i = \bar{P}_i\, \bar{\boldsymbol{s}}_i + \hat{P}_i\, \hat{\boldsymbol{s}}_i,$$

where $\mathrm{q} \times |\bar{\boldsymbol{\mu}}_i|$ matrix $\bar{R}_i$ and $\mathrm{q} \times |\hat{\boldsymbol{\mu}}_i|$ matrix $\hat{R}_i$ consist of those columns of $R^T$ that are selected by $\bar{\boldsymbol{\mu}}_i$ and $\hat{\boldsymbol{\mu}}_i$, respectively. Analogously, $\mathrm{q} \times |\bar{\boldsymbol{\nu}}_i|$ matrix $\bar{P}_i$ and $\mathrm{q} \times |\hat{\boldsymbol{\nu}}_i|$ matrix $\hat{P}_i$ are formed from the columns of $P^T$ selected by $\bar{\boldsymbol{\nu}}_i$ and $\hat{\boldsymbol{\nu}}_i$, respectively. These vector and matrix partitions are illustrated in Figure S1C. Introducing abbreviations

$$\boldsymbol{x}_i = \begin{bmatrix} \hat{\boldsymbol{j}}_i \\ \hat{\boldsymbol{s}}_i \end{bmatrix} \quad \text{and} \quad \boldsymbol{k}_i = \begin{bmatrix} \bar{R}_i & \bar{P}_i \end{bmatrix} \begin{bmatrix} \bar{\boldsymbol{j}}_i \\ \bar{\boldsymbol{s}}_i \end{bmatrix},$$

an equivalent reformulation of Equation 7 reads

$$\begin{bmatrix} \hat{R}_i & \hat{P}_i \end{bmatrix} \boldsymbol{x}_i = -\boldsymbol{k}_i, \quad i = 1, 2, \ldots \mathrm{n}. \tag{8}$$

The point of such algebraic acrobatics is that Equation 8 represents systems of linear equations, each in the

$$\mathrm{u}_i = |\hat{\boldsymbol{\mu}}_i| + |\hat{\boldsymbol{\nu}}_i|$$

unknown parameters $\boldsymbol{x}_i$, compared to Equation 7 in which the solution vector comprised unknown and known components. It thus allows to study the identifiability of $\boldsymbol{x}_i$.

### Identifiability conditions

Clearly, Equation 8 is underdetermined if

$$\mathrm{d}_i = \mathrm{u}_i - \operatorname{rank}(\begin{bmatrix} \hat{R}_i & \hat{P}_i \end{bmatrix}) > 0.$$

To analyse this solution space dimensionality, let $\mathrm{n} \times |\hat{\boldsymbol{\mu}}_i|$ matrix $\hat{J}_i^{-1}$ consist of the columns of $\left(J^{-1}\right)^T$ that are selected by $\boldsymbol{\mu}_i$. Similarly, $|\bar{\boldsymbol{\nu}}_i| \times \mathrm{n}$ matrix $\bar{S}^i$ and $\hat{\boldsymbol{\nu}}_i \times \mathrm{n}$ matrix $\hat{S}^i$ shall be formed by taking rows of $S^T$ according to indices in $\bar{\boldsymbol{\nu}}_i$ and $\hat{\boldsymbol{\nu}}_i$, as shown in Figure S1D. Also, we have $I_i$ denote the $i$-dimensional identity matrix and $0_{i,j}$ the $i \times j$ zero-matrix. We use these definitions and Equation 4 to write

$$\hat{R}_i = -P^T S^T \hat{J}_i^{-1} \quad \text{and} \quad P^T S^T = \begin{bmatrix} \hat{P}_i & \bar{P}_i \end{bmatrix} \begin{bmatrix} \hat{S}_i \\ \bar{S}_i \end{bmatrix},$$

and arrive at

$$\begin{bmatrix} \hat{R}_i & \hat{P}_i \end{bmatrix} = -\begin{bmatrix} \hat{P}_i & \bar{P}_i \end{bmatrix} \Psi_i, \quad \text{with} \tag{9}$$

$$\Psi_i = \begin{bmatrix} \hat{S}_i \hat{J}_i^{-1} & I_{|\hat{\boldsymbol{\nu}}_i|} \\ \bar{S}_i \hat{J}_i^{-1} & 0_{|\bar{\boldsymbol{\nu}}_i|,|\hat{\boldsymbol{\nu}}_i|} \end{bmatrix}. \tag{10}$$

Note that $\begin{bmatrix} \hat{P}_i & \bar{P}_i \end{bmatrix}$ is nothing but a rearrangement of the columns of $P^T$ and therefore

$$\operatorname{rank}(\begin{bmatrix} \hat{P}_i & \bar{P}_i \end{bmatrix}) = \operatorname{rank}(P) = \mathrm{p}.$$

Claiming $P$ to have rank p assumes that throughout the experiment every type of perturbation was applied in a non-trivial combination. This is not a limiting constraint as it is for example satisfied for a perturbation scheme in which each type of perturbation is applied once individually, which is the case for the examples discussed here.

From $\begin{bmatrix} \hat{P}_i & \bar{P}_i \end{bmatrix}$ having full (column) rank follows that

$$\begin{aligned} &\operatorname{rank}([\hat{R}_i \quad \hat{P}_i]) = \operatorname{rank}(\Psi_i) \\ &= \operatorname{rank}([\hat{S}_i \hat{J}_i^{-1} \quad I_{|\hat{\boldsymbol{\nu}}_i|}]) + \operatorname{rank}([\bar{S}_i \hat{J}_i^{-1} \quad 0_{|\bar{\boldsymbol{\nu}}_i|,|\hat{\boldsymbol{\nu}}_i|}]) \\ &= |\hat{\boldsymbol{\nu}}_i| + \operatorname{rank}\left(\bar{S}_i \hat{J}_i^{-1}\right), \end{aligned}$$

so that the solution subspace has dimensionality

$$\mathrm{d}_i = |\hat{\boldsymbol{\mu}}_i| - \operatorname{rank}\left(\bar{S}_i \hat{J}_i^{-1}\right).$$

From the dimensionality of matrix product $\bar{S}_i \hat{J}_i^{-1}$ we can conclude that $d_i \geq \max(0, \mathrm{n} - |\bar{\boldsymbol{\mu}}_i| - |\bar{\boldsymbol{\nu}}_i|)$. Thus, to fully determine $\boldsymbol{x}_i$ we need to provide at least as many elements of prior knowledge as there are nodes in the network, which agrees with our earlier observation that we can transform the rate equations with an arbitrary $\mathrm{n} \times \mathrm{n}$ matrix without altering the steady states.

If indeed $\mathrm{d}_i > 0$, there is a $\mathrm{u}_i \times \mathrm{d}_i$ matrix $V_i$ whose columns form a basis of the kernel of $\begin{bmatrix} \hat{R}_i & \hat{P}_i \end{bmatrix}$, so that, given $\tilde{\boldsymbol{x}}_i$, a specific solution to Equation 8, any

$$\boldsymbol{x}_i = V_i \boldsymbol{w} + \tilde{\boldsymbol{x}}_i, \ \forall \boldsymbol{w} \in \mathbb{R}^{\mathrm{d}_i} \tag{11}$$

is also a solution of Equation 8. But even though the equation system is then underdetermined, not all network parameters are necessarily unidentifiable. Rather,

$$\begin{aligned} [\boldsymbol{x}_i]_j \text{ identifiable} &\iff \boldsymbol{e}_j^T V_i = 0 \\ &\iff \exists\, \boldsymbol{w} \in \mathbb{R}^{\mathrm{q}} : \begin{bmatrix} \hat{R}_i & \hat{P}_i \end{bmatrix}^T \boldsymbol{w} = \boldsymbol{e}_j, \end{aligned} \tag{12}$$

where $\boldsymbol{e}_j$ is the $j$-th standard basis vector of according length. We shall use Equation 9 to reformulate this identifiability condition. To this end, recall the earlier assertion about the full (column) rank of $\begin{bmatrix}\hat{P}_i & \bar{P}_i\end{bmatrix}$, from which follows that

$$\forall\, \tilde{\boldsymbol{w}} \in \mathbb{R}^{\mathrm{p}},\ \exists\, \boldsymbol{w} \in \mathbb{R}^{\mathrm{q}} :\ \tilde{\boldsymbol{w}}^T = \boldsymbol{w}^T \begin{bmatrix}\hat{P}_i & \bar{P}_i\end{bmatrix},$$

so that we can write

$$[\boldsymbol{x}_i]_j \text{ identifiable} \iff \exists\, \tilde{\boldsymbol{w}} \in \mathbb{R}^{\mathrm{p}} : \tilde{\boldsymbol{w}}^T\, \Psi_i = \boldsymbol{e}_j^T .$$

Next, let $\tilde{\boldsymbol{w}}_1$ and $\tilde{\boldsymbol{w}}_2$ consist of the first $|\hat{\boldsymbol{\nu}}_i|$ and the last $|\bar{\boldsymbol{\nu}}_i|$ components of $\tilde{\boldsymbol{w}}$, such that $\tilde{\boldsymbol{w}}^T = [\tilde{\boldsymbol{w}}_1^T \quad \tilde{\boldsymbol{w}}_2^T]$. Accordingly, standard base vector $\boldsymbol{e}_j$ is split into its first $|\hat{\boldsymbol{\mu}}_i|$ and last $|\hat{\boldsymbol{\nu}}_i|$ components, $\boldsymbol{e}_j^T = [\boldsymbol{f}_j^T \quad \boldsymbol{g}_j^T]$. This allows to rewrite the previous equation as

$$\tilde{\boldsymbol{w}}_1 = \boldsymbol{g}_j, \text{ and}$$
$$\tilde{\boldsymbol{w}}_2^T \left(\bar{S}_i \hat{J}_i^{-1}\right) = \boldsymbol{f}_j^T - \boldsymbol{g}_j^T \left(\hat{S}_i \hat{J}_i^{-1}\right).$$

Recall that $[x_i]_j$ denotes unknown interaction strengths for $j \leq |\hat{\boldsymbol{\nu}}_i| \iff \boldsymbol{g}_j = \boldsymbol{0}$ and thus

$$[\hat{\boldsymbol{j}}_i]_j \text{ identifiable} \iff \operatorname{rank}\left(\begin{bmatrix}\bar{S}_i \hat{J}_i^{-1} \\ \boldsymbol{f}_j^T\end{bmatrix}\right) = \operatorname{rank}(\bar{S}_i \hat{J}_i^{-1})$$
$$\iff 1 + \operatorname{rank}(\bar{S}_i \hat{J}_{i\setminus j}^{-1}) = \operatorname{rank}(\bar{S}_i \hat{J}_i^{-1}), \tag{13}$$

where $\hat{J}_{i\setminus j}^{-1}$ is matrix $\hat{J}_i^{-1}$ with the $j$-th column removed. For the unknown sensitivity coefficients, where $j > |\hat{\boldsymbol{\nu}}_i| \iff \boldsymbol{f}_j = \boldsymbol{0}$, we find the identifiability conditions

$$[\hat{\boldsymbol{s}}_i]_j \text{ identifiable}$$
$$\iff \operatorname{rank}\left(\begin{bmatrix}\bar{S}_i \\ \hat{\boldsymbol{S}}_i^j\end{bmatrix} \hat{J}_i^{-1}\right) = \operatorname{rank}(\bar{S}_i \hat{J}_i^{-1}), \tag{14}$$

where $\hat{\boldsymbol{S}}_i^j$ denotes the $j$-th row of matrix $\hat{S}_i$.

### Structural identifiability

The identifiability conditions in equations 13 and 14 relate the identifiability of the unknown parameters to a discussion of the rank of matrix product $\bar{S}_i \hat{J}_i^{-1}$. The product however depends on the unknown parameters themselves, so that its rank cannot be directly computed. Here we show that a reasonable assumption make this possible nevertheless and allows to express the identifiability conditions as a very intuitive maximum flow problem.

First, we rewrite the identity $J^{-1}J = I_{\mathrm{n}}$ as

$$[J^{-1}]_{kl} = \sum_{m \neq l} [J^{-1}]_{km}\, [J]_{ml} - \delta_{kl},$$

with $\delta_{kl}$ being the Kronecker delta (recall that $J_{ll} = -1$). We can view this equation as a recurrence relation and repeatedly replace the $[J^{-1}]_{km}$ terms in the sum. The sum contains non-vanishing terms for each edge that leaves node $l$. Therefore, each replacement leads to the next downstream node, so that eventually one arrives at

$$[J^{-1}]_{kl} = l \rightarrowtail k\ \ [J^{-1}]_{kk}, \text{ with}$$
$$l \rightarrowtail k = \sum_{\boldsymbol{\omega} \in \Omega_{l \to k}} \prod_{m=1}^{|\boldsymbol{\omega}|-1} [J]_{\omega_{m+1}\, \omega_m},$$

where the set $\Omega_{l\to k}$ contains elements, $\boldsymbol{\omega}$, for every path from node $l$ to node $k$, each of which lists the nodes along that path. Strictly speaking, these elements are walks rather than paths because some nodes will appear multiple times if loops exist between $l$ and $k$. With loops, $\Omega_{l\to k}$ even contains an infinite number of walks of unbounded lengths. But as the real part of all eigenvalues of $J$ are assumed negative, the associated products of interaction strengths will eventually converge to zero with increasing walk length. Here however, we can safely ignore these subtleties.

To simplify our notation, we want to expand the network by considering perturbations $\bar{\boldsymbol{\nu}}_i$ as additional nodes, each with edges that are directed towards that perturbation's targets. Furthermore, letting the interaction strength associated with these new edges be given by the appropriate entries in $S$ we can rewrite the matrix product

$$\left[\bar{S}_i \hat{J}_i^{-1}\right]_{kl} = \bar{\nu}_{ik} \rightarrowtail \hat{\mu}_{il}\ [J^{-1}]_{\hat{\mu}_{il}\, \hat{\mu}_{il}}$$

where $\hat{\mu}_{il}$ and $\bar{\nu}_{il}$ denote the $l$-th entry in $\hat{\boldsymbol{\mu}}_i$ and $\bar{\boldsymbol{\nu}}_i$, respectively. As every finite-dimensional matrix has a rank decomposition, we can further write

$$\bar{S}_i \hat{J}_i^{-1} = \Upsilon_i\, Y_i, \tag{15}$$

where $|\bar{\boldsymbol{\nu}}_i| \times \operatorname{rank}(\bar{S}_i \hat{J}_i^{-1})$ matrix $\Upsilon_i$ and $\operatorname{rank}(\bar{S}_i \hat{J}_i^{-1}) \times |\hat{\boldsymbol{\mu}}_i|$ matrix $Y_i$ have full rank. Finding such a decomposition therefore reveals the rank of $\bar{S}_i \hat{J}_i^{-1}$. To this end, we propose

$$[\Upsilon_i]_{kn} = \bar{\nu}_{ik} \rightarrowtail y_{in}, \text{ and } [Y_i]_{nl} = y_{in} \rightarrowtail \hat{\mu}_{il}\ [J^{-1}]_{\hat{\mu}_{il}\, \hat{\mu}_{il}},$$

where $y_{in}$ denotes the $n$-th component of a certain node set $\boldsymbol{y}_i$. In order for Equation 15 to hold, it must be possible to split each path from any perturbation $\bar{\nu}_{il}$ to any node $\hat{\mu}_{il}$ into a section that leads from the perturbation to a node in $\boldsymbol{y}_i$ and a subsequent section that leads from this node to $\hat{\mu}_{il}$. For an extended graph that includes an additional source node, with outgoing edges to each perturbation in $\bar{\boldsymbol{\nu}}_i$, and an additional sink node, with incoming edges from all nodes in $\hat{\boldsymbol{\mu}}_i$ (see Figure S2A), $\boldsymbol{y}_i$ thus constitutes a vertex cut whose removal disconnects the graph and separates the source and the sink node into distinct connected components. Next, we want to show that if $\boldsymbol{y}_i$ is a minimum vertex cut, the rank of $\bar{S}_i \hat{J}_i^{-1}$ equals the size of $\boldsymbol{y}_i$. Because Equation 15 is a rank decomposition this is equivalent to showing that the according matrices $\Upsilon_i$ and $Y_i$ have full rank. To do so we apply Menger's theorem (Menger, 1927), which states that the minimal size of $\boldsymbol{y}_i$ equals the maximum number of vertex-disjoint paths from the source to the sink node. This also implies that each of these vertex-disjoint paths goes through a different node of the vertex cut $\boldsymbol{y}_i$. Recall that entries in $\Upsilon_i$ constitute sums over paths from perturbation to vertex cut nodes, so that we could write

$$\Upsilon_i = \bar{\Upsilon}_i + \hat{\Upsilon}_i,$$

where $\bar{\Upsilon}_i$ only contains the vertex-disjoint paths and $\hat{\Upsilon}_i$ the sums over the remaining paths. As each of these vertex disjoint paths ends in a different vertex cut node, any column in $\bar{\Upsilon}_i$ can contain no more than a single non-zero entry. Furthermore, as a consequence of Menger's theorem there are exactly $|\boldsymbol{y}_i|$ non-zero columns. Because these paths are indeed vertex disjoint also no row in $\bar{\Upsilon}_i$ has more than a single non-zero entry. Thus, the non-zero columns are independent, showing that $\bar{\Upsilon}_i$ has full rank. The crucial assumption we want to make now is that the values of the interaction strengths lie outside a specific algebraic variety, which would render $\bar{\Upsilon}_i + \hat{\Upsilon}_i$ rank deficient. This would for example be the case if for a given vertex disjoint path there also is an alternative path whose associated product of interaction strengths has the same magnitude as that of the vertex disjoint path but opposite sign, making their sum vanish. This effect corresponds to a perfectly self-compensating perturbation. Most biological networks cannot fine-tune their interactions to such a degree that they could achieve perfect

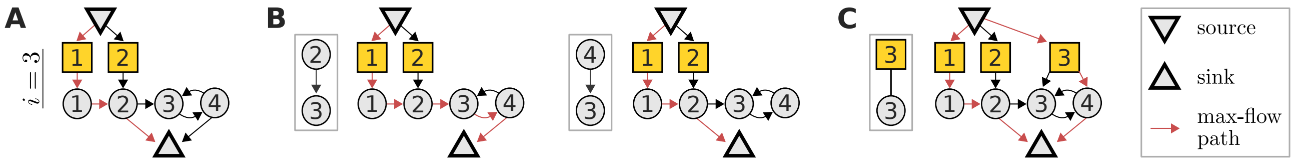


**Figure S2:** A maximum flow problem determines the identifiability of interaction strengths and perturbation sensitivities when reconstructing a network from perturbation data. Here, this is illustrated for the toy model from Figure S1A. To inquire about the identifiability of either edges going into node 3, or the sensitivity of node 3 to perturbations, we construct a flow network (**A**) with unit edge and node capacities, as described in the text. We highlight in red a path carrying the maximal flow of one. While this max-flow path is not unique, no other combination of paths could yield a larger flow. The interaction strength between a given node and node 3 is identifiable if and only if the maximum flow is reduced after removing that node's edge to the sink node (**B**). Yet here, we can always find alternative max-flow paths that re-establish a unit-flow after removal of the according edges. Thus the respective edges are non-identifiable. Similarly, the sensitivity of node 3 to perturbation 3 is identifiable if and only if a specific extension of the flow network (**C**) does not increase the maximum flow. But here the maximum flow is indeed increased by one, which again reveals non-identifiability. Such flow representations also provide an intuitive understanding on how alterations in the network or perturbation setting affect identifiability. For example, it is obvious that if the toy model would not contain an edge from node 3 to 4, the edge from 2 to 3 would become identifiable.

self-compensation, which justifies the non-cancellation assumption.

The same line of reasoning will demonstrate a full rank for matrix $Y_i$ as well, which implies that indeed

$$\operatorname{rank}(\bar{S}_i \hat{J}_i^{-1}) = |\boldsymbol{y}_i|,$$

where $\boldsymbol{y}_i$ is a minimum vertex cut between source and sink node. This equation has the crucial benefit that $|\boldsymbol{y}_i|$ does not depend on unknown matrix entries. Rather, the size of the minimum vertex cut is equal to the defined maximum flow from source to sink node with all nodes having unit capacity (Ahuja *et al.*, 1993), as detailed in Figure S2A. The resulting maximum flow problem can be solved in only $\mathcal{O}\left(\mathrm{n}^{1/2}\sum_i^{\mathrm{n}}(|\hat{\boldsymbol{\mu}}_i| + |\hat{\boldsymbol{\nu}}_i|)\right)$, as shown in Theorem 6.3 in (Even and Even, 2012). More importantly though, it allows to express the algebraic identifiability conditions in terms of network properties and thus allows to intuitively understand the relationship between network topology, perturbation targets and identifiability. To clarify this, recall Equation 13, the condition for the identifiability of network edges $\hat{\boldsymbol{\mu}}_i$. We can now restate it as follows. Interaction strength $[\hat{\boldsymbol{j}}_i]_j$ is identifiable if and only if the removal of the edge from node $\hat{\mu}_{ij}$ to the sink node reduces the maximum flow of the network, see Figure S2B. Similarly, the condition for identifiability of sensitivitiy $[\hat{\boldsymbol{s}}_i]_j$, as expressed by Equation 14 is fulfilled when the maximum flow does not increase when an additional edge connects the source node with perturbation node $\hat{\nu}_{ij}$, see Figure S2B.

In summary, a reasonable non-cancellation assumption allows to map the analysis of identifiability from an algebraic description, that required a specification of unknown matrix entries, to a maximum-flow problem that only depends on the network topology and perturbation targets. In addition, such a graphic representation provides a overview on how network and perturbation modifications relate to changes in identifiability, as explained in Figure S2. This enables a straightforward optimization of experimental design, as discussed further below.

### S2 Identifiability relationships

Network inference typically is an underdetermined problem for which the number of measurements falls short on the number of unknown interaction terms (De Smet and Marchal, 2010; Gross *et al.*, 2019), resulting in many non-identifiable parameters. To tackle this problem, we could construct identifiable models by fixing certain parameters to some constant values. Clearly, the remaining, inferred parameter values will then disagree with those that would have been obtained from a fully-determining experiment. Nevertheless, such effective models are useful as they allow for meaningful comparisons of the inferred parameters between perturbation experiments on similar systems, e.g. when studying the same signalling pathway in different cell lines (Bosdriesz *et al.*, 2018). To derive such a determined system requires to study the relationship between non-identifiable parameters in the sense that we ask which parameters need to be fixed in order to render which other parameters identifiable. Even though the dimensionality of the solution space, $\mathrm{d}_i$, is known, this question is not trivial, because even groups with $\mathrm{d}_i$ or fewer parameters might already be linearly dependent and fixing them will therefore not effectively reduce the degrees of freedom of the equation system.

Take as example a case where the first two rows of kernel matrix $V_i$ from Equation 11, are linearly dependent, that is $\alpha\, \boldsymbol{V}_i^1 = \boldsymbol{V}_i^2$. Then $[\boldsymbol{x}_i]_1$ and $[\boldsymbol{x}_i]_2$ are linearly dependent as well, $[\boldsymbol{x}_i]_2 = \boldsymbol{V}_i^2\, \boldsymbol{v} = \alpha\, \boldsymbol{V}_i^1\, \boldsymbol{v} = \alpha\, [\boldsymbol{x}_i]_1$, which implies that $[\boldsymbol{x}_i]_2$ becomes identifiable if $[\boldsymbol{x}_i]_1$ is known, and vice versa, even if $\mathrm{d}_i > 1$ ($\tilde{\boldsymbol{x}}_i$ was dropped to simplify notation). Moreover, prior knowledge on both $[\boldsymbol{x}_i]_1$ and $[\boldsymbol{x}_i]_2$ would overdetermine this linear subsystem and not further reduce the degrees of freedom for the remaining unknown parameters. Examining such parameter dependencies is a direct generalization of the original identifiability condition in Equation 12. There, identifiability of an unknown parameter relied on a $V_i$-row being zero, that is, on a one-row submatrix being rank deficient. Now, we inspect not only single but groups of $V_i$-rows for rank deficiency. But which groups of rows should we consider to achieve an effective description of dependency? To answer this question let us first generalize the previous example.

We were asking if the $j$-th $\boldsymbol{x}_i$ component becomes identifiable if a set of other $\boldsymbol{x}_i$ components is known. With $\mathcal{I}$ denoting the set of indices of these other components, let us recall Equation 11 and name their homogenous parts

$$\hat{x}_i^{\mathcal{I}} = \boldsymbol{V}_i^j\, \boldsymbol{v} \quad \text{and} \quad \bar{\boldsymbol{x}}_i^{\mathcal{I}} = V_i^{\mathcal{I}}\, \boldsymbol{v},$$

where $\boldsymbol{V}_i^j$ is the $j$-th row of $V_i$, and $V_i^{\mathcal{I}}$ the matrix that gathers all $V_i$ rows with indices in $\mathcal{I}$. We can then put down a formal identifiability statement

$$\begin{aligned} &\exists\, \mathcal{I} \subseteq \{1, \ldots, \mathrm{u}_i\} \setminus j,\ \exists\, \boldsymbol{w} \in \mathbb{R}^{|\mathcal{I}|} :\ \boldsymbol{V}_i^j = \boldsymbol{w}^T\, V_i^{\mathcal{I}} \\ &\iff \hat{x}_i^{\mathcal{I}} = \boldsymbol{w}^T\, V_i^{\mathcal{I}}\, \boldsymbol{v} = \boldsymbol{w}^T \bar{\boldsymbol{x}}_i^{\mathcal{I}}. \end{aligned} \tag{16}$$

In other words, if the $j$-th $V_i$-row lies within the row-space of the set of $V_i$-rows with indices $\mathcal{I}$, the $j$-th unknown parameter can be expressed as a linear combination of the set of parameters with indices $\mathcal{I}$. This means that knowledge of the set of parameters with indices $\mathcal{I}$ then implies identifiability of the $j$-th parameter. However, this statement does not imply the uniqueness of $\mathcal{I}$. On the contrary, if the $j$-th $V_i$-row lies within the $\mathcal{I}$-associated rowspace, it will also do so if additional $V_i$ rows are added to the set. Similarly, there could be a linearly dependent subset of $V_i$-rows that all lie within the $\mathcal{I}$ associated row-space. This would allow for multiple row-combinations to span the $\mathcal{I}$-associated rowspace and thus implicate the identifiability statement. Both cases show, that various combinations of additionally fixed parameters can imply the identifiability of a certain other parameter.

A comprehensive description of this combinatorial space arises from a mathematical structure that has been termed matroid (Whitney, 1935). Matroids are a generalized description of linear independence in vector spaces. Here we are concerned with representable matroids, which are those that specify linear (in-)dependence of any combination of columns of a matrix. Amongst their various equivalent definitions, the one that relates directly to our problem is the definition in terms of cyclic flats (also called circuit closures) and their ranks (Oxley, 2006). To specify these we need to define a few terms. First, let $\mathcal{E}$ be the ground set of matroid $\mathcal{M}$, that is, the set of indices enumerating the columns of the associated matrix. Furthermore, define a circuit as a dependent set (of columns) whose proper subsets are all independent. The set of circuits can be enumerated with an incremental polynomial-time algorithm (Boros *et al.*, 2003). Finally, we define a flat as a subset of $\mathcal{E}$, with the associated submatrix having rank $r$, such the addition of any other element to the set would increase the rank. With this we can define $\mathcal{C}_r$, a cyclic flat of rank $r$, as a flat that is the union of a set of circuits with rank $r$. We show in the next section how to obtain cyclic flats from circuits and vice versa.

Let us now consider $\mathcal{M}_i$, the matroid whose groundset $\varepsilon_i$ covers the $\mathrm{u}_i$ columns of $(V_i)^T$. Each element in $\varepsilon_i$ is thus associated with an unknown parameter. The key inside is that $\mathcal{M}_i$'s set of circuits fully characterizes the identifiability relationships between the non-identifiable parameters. This is because the circuit dependency implies that any parameter represented by a given circuit element is identifiable when the remaining circuit elements are known. Additionally, this set of remaining parameters is guaranteed to be minimal because they are linearly independent. The enumeration of the circuits with the aforementioned algorithm requires a dependence oracle that indicates whether a column subset is dependent or not. For this, we first consider another matroid $\mathcal{M}'_i$, which is associated with the $\mathrm{u}_i$ columns of $\Psi_i$, as defined in Equation 10. Because $V_i$ spans the kernel of matrix $\Psi_i$, $\mathcal{M}'_i$ is dual to $\mathcal{M}_i$ (Whitney, 1935). This implicates that the rank of the $(V_i)^T$ column-subset $\mathcal{I}$ relates to that of the complementary columns $\tilde{\mathcal{I}} = \varepsilon_i \setminus \mathcal{I}$ of $\Psi_i$ as follows

$$\mathrm{rank}_{\mathcal{M}_i}(\mathcal{I}) = \mathrm{rank}_{\mathcal{M}'_i}(\tilde{\mathcal{I}}) + |\mathcal{I}| - (\mathrm{u}_i - \mathrm{d}_i).$$

To investigate the dual rank, we note that we can establish the column subset of $\Psi_i$ by a right multiplication with the $\mathrm{u}_i \times |\tilde{\mathcal{I}}|$ matrix $\mathcal{P}$, which is an identity matrix where columns that correspond to missing indices in $\tilde{\mathcal{I}}$ are removed. Furthermore, we subdivide elements in $\tilde{\mathcal{I}}$ into sets $\tilde{\mathcal{I}}_1$ and $\tilde{\mathcal{I}}_2$ based on whether they are less than or equal to $|\hat{\boldsymbol{\mu}}_i|$ or not, which allows to define matrices $\mathcal{P}_1$ and $\mathcal{P}_2$ by the partitioning

$$\mathcal{P} = \begin{bmatrix} \mathcal{P}_1 & 0_{|\hat{\boldsymbol{\mu}}_i|,|\tilde{\mathcal{I}}_2|} \\ 0_{|\hat{\boldsymbol{\nu}}_i|,|\tilde{\mathcal{I}}_1|} & \mathcal{P}_2 \end{bmatrix}. \tag{17}$$

Then,

$$\begin{aligned} \mathrm{rank}_{\mathcal{M}'_i}(\tilde{\mathcal{I}}) &= \mathrm{rank}(\Psi_i \mathcal{P}) \\ &= \mathrm{rank}\left(\begin{bmatrix} \hat{S}_i \hat{J}_i^{-1} \mathcal{P}_1 & \mathcal{P}_2 \\ \check{S}_i \hat{J}_i^{-1} \mathcal{P}_1 & 0_{|\check{\boldsymbol{\nu}}_i|,|\tilde{\mathcal{I}}_2|} \end{bmatrix}\right) \\ &= |\tilde{\mathcal{I}}_2| + \mathrm{rank}\left(\begin{bmatrix} \tilde{\mathcal{P}}_2^T \hat{S}_i \\ \check{S}_i \end{bmatrix} \hat{J}_i^{-1} \mathcal{P}_1\right), \end{aligned} \tag{18}$$

where $\tilde{\mathcal{P}}_2$ is the identity matrix without the columns that appear in $\mathcal{P}_2$. Left-multiplication by $\tilde{\mathcal{P}}_2^T$ thus selects rows that correspond to missing indices in $\tilde{\mathcal{I}}_2$. The crucial point of this calculation is that we arrived at a matrix product that has the same form as the one discussed in the previous section. Therefore, the dual rank can be evaluated independently of the unknown entries in $J$ and $S$ because the last term in the previous equation equals to the maximum flow through the associated network, with connections from the source and to the sink nodes that are chosen according to $\tilde{\mathcal{I}}$, as shown. This allows to construct the oracle and identify the set of circuits. Therefore the identifiability relationships between unknown parameters can be inferred from information about network topology and perturbation targets alone.

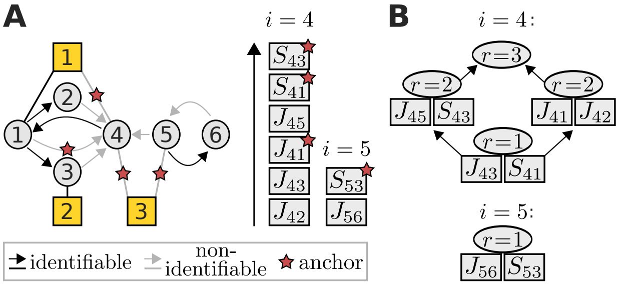


**Figure S3:** In this toy network (**A**), nodes 4 and 5 are associated with non-identifiable parameters. These can take values from certain linear sub-spaces whose hierarchy is represented by the lattices of cyclic flats of rank $r$ (**B**). Each cyclic flat consists of the annotated elements in addition to elements from its preceding cyclic flats. To achieve identifiability requires to set certain parameters to a constant value. A preference to which parameters this should be is represented here as a ranked list (arrow indicates direction of increasing preference). The matroid formalism identifies the smallest and most preferred set of parameters that, when set to a constant value, render the network model fully identifiable. Here these are marked by red stars.

Instead of listing the set of circuits, we propose cyclic flats as an equivalent but more concise representation of the identifiability relationships. They form a geometric lattice when ordered by inclusion (a cyclic flat precedes another if it is its proper subset) and can thus be graphically represented as a compact hierarchical structure. We demonstrate this for the example network shown in Figure S3. The depicted lattice makes the identifiability relationships evident. All elements of a cyclic flat with rank $r$ become identifiable if at least $r$ independent flat elements are fixed. A set of elements is independent if fixing any combination if its elements does not render any of its other element identifiable. Let us clarify this at an example where we are interested in determining the parameters that need to be fixed in order to make $J_{43}$ identifiable. Following the previous rules, Figure S3 reveals that this could be achieved by fixing either $S_{41}$ alone, or the parameter pairs $J_{45} \cup S_{43}$ or $J_{41} \cup S_{42}$. In the latter two cases $S_{53}$ would become identifiable as well.

When the goal is to achieve a fully identifiable network model, as discussed before, there typically are preferences as to which non-identifiable parameters should be fixed. For example, if there is noisy external data on parameter values we would rather fix those parameters values in which we have high confidence. Or, if we are to construct the aforementioned effective signalling models for the comparison of different cell lines, we would want to fix those parameters, which we expect to be equal between different cell lines and infer those parameters for which cell line differences are expected (Bosdriesz *et al.*, 2018). Thus, in these scenarios fixing of each parameter is associated with a certain preference (weight) and our goal is to find a minimum number of parameters that need to be fixed such that their sum of weights is maximal. In fact, matroids owe their striking appearance in combinatorial optimization because this problem is solvable with the Greedy Algorithm (Oxley, 2003; Gale, 1968): Amongst the set of non-identifiable parameters in

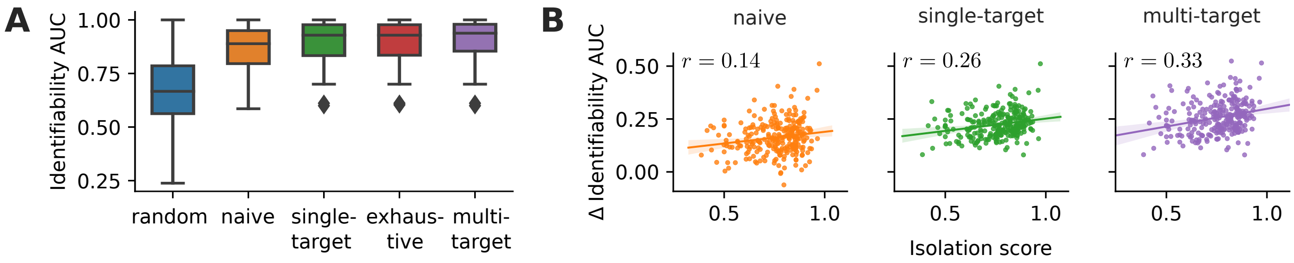


**Figure S4:** Performance of different experimental design strategies on 78 human KEGG pathways with up to 15 nodes (**A**). The single-target and exhaustive strategies show identical performance. The difference between the identifiability AUC between non-random to random strategies negatively correlates (Spearman correlation coefficient $r$) with the isolation score (**B**). Here, all human KEGG pathways are considered.

$\epsilon_i$, sequentially select the parameters with highest weight, that have not yet become identifiable from fixing the so-far selected set. Thus, instead of providing numerical weights for unknown parameters it is sufficient to rank them. We depict examples of such ordered lists in Figure S3 and show the resulting fully identifiable maximum-weight-model.

#### Circuits and circuit closures

As both, the set of circuits and the circuit closures combined with their ranks, are an equivalent definition of a matroid they imply each other. Recall that circuits that contain a given network parameter describe the minimal sets of network parameters that need to be fixed to render that parameter identifiable. The flat of closures conveniently display these circuits as follows. By definition, any circuit is a $r+1$-element subset, $\mathcal{S}$, of some cyclic flat $\mathcal{C}_r$ with rank $r$. Thus, to obtain all circuits containing a certain parameter, consider all such subsets of cyclic flats that include this parameter. Yet $\mathcal{S}$ is only a circuit if none of its subsets $\underline{\mathcal{S}} \subset \mathcal{S}$ is dependent, in which case there is another circuit $\underline{\mathcal{C}} \subseteq \underline{\mathcal{S}}$. Since the lattice of cyclic flats is ordered by inclusion, $\underline{\mathcal{C}}$ is a subset of a cyclic flat that precedes $\mathcal{C}_r$ in the lattice. Therefore, $\mathcal{S}$ is only a circuit if no cyclic flat preceding to $\mathcal{C}_r$ contains a circuit that is a proper subset of $\mathcal{S}$.

We mentioned that circuits can be enumerated in incremental polynomial-time (Boros *et al.*, 2003). In a next step, we generated circuit closures from the set of circuits. To this end, we first order circuits by size and iterate through that list. For each circuit of a given rank we identify circuits of up to its size whose intersection is equal or larger to its rank. Their union forms a circuit closure. Next, one continues the circuit iteration while skipping circuits that have already been assigned to a circuit closure. Eventually, this generates the entire ensemble of circuit closures. Find an implementation in the function `circuits2cyclic_flats` which is part of the `identifiability` module of the IdentiFlow package available at `github.com/GrossTor/IdentiFlow`.

### S3 Perturbation experiments for KEGG pathways

KEGG data (Kanehisa *et al.*, 2019) was retrieved using the KEGG API. We retrieved KGML files for human pathways and from them build network representations based on their 'relation elements'. For each such representation we computed the size of its largest connected component. The pathway was filtered out if it was smaller than five.

The performance of the exhaustive strategy could be observed for small pathways Figure S4 A. In addition, we further confirmed our hypothesis that the isolation score is predictive with respect to the performance of the design strategies Figure S4 B.
